# Improved inference of latent neural states from calcium imaging data

**DOI:** 10.1101/2025.10.17.682993

**Authors:** Stephen Keeley, David Zoltowski, Adam Charles, Jonathan Pillow

## Abstract

Calcium imaging (CI) is a standard method for recording neural population activity, as it enables simultaneous recording of hundreds-to-thousands of individual somatic signals. Accordingly, CI recordings are prime candidates for population-level latent variable analyses, for example, using models such as Gaussian Process Factor Analysis (GPFA), hidden Markov models (HMMs), and latent dynamical systems. However, these models have been primarily developed and fine-tuned for electrophysiological measurements of spiking activity. To adapt these models for use with the calcium signals recorded with CI, per-neuron fluorescence time-traces are typically either de-convolved to approximate spiking events or analyzed directly under Gaussian observation assumptions. The former approach, while enabling the direct application of latent variable methods developed for spiking data, suffers from the imprecise nature of spike estimation from CI. Moreover, isolated spikes can be undetectable in the fluorescence signal, creating additional uncertainty. A more direct model linking observed fluorescence to latent variables would account for these sources of uncertainty. Here, we develop accurate and tractable models for characterizing the latent structure of neural population activity from CI data. We propose to augment HMM, GPFA, and dynamical systems models with a CI observation model that consists of latent Poisson spiking and autoregressive calcium dynamics. Importantly, this model is both more flexible and directly compatible with standard methods for fitting latent models of neural dynamics. We demonstrate that using this more accurate CI observation model improves latent variable inference and model fitting on both CI observations generated using state-of-the-art biophysical simulations and imaging data recorded in an experimental setting. We expect the developed methods to be widely applicable to many different analyses of population CI data.

## Introduction

Electrophysiological recordings have historically been the de-facto approach for observing the activity of single neurons. Consequently, many statistical models for neuronal data at the single-cell level are formulated for spike train observations. That is, they describe a mapping from an observed covariate or unobserved latent variables to a probability distribution over discrete spike events. This class of models includes (but is not limited to) Poisson regression models (***Truccolo et al., 2005***; ***Pillow et al., 2008***; ***McFarland et al., 2013***; ***Park et al., 2013***; ***Zoltowski and Pillow, 2018***; ***Medvedeva et al., 2026***), non-Poisson spike count regression models (***Pillow and Scott, 2012***; ***Goris et al., 2014***; ***Williamson et al., 2015***; ***Gao et al., 2015***; ***Linderman et al., 2016a***; ***Stevenson, 2016***; ***Charles et al., 2018***), linear dynamical systems (LDS) (***Macke et al., 2011***), nonlinear dynamics models (***Linderman et al., 2017***; ***Pandarinath et al., 2018***; ***Duncker et al., 2019***; ***Zhao and Park, 2019***), Gaussian process factor analysis (GPFA) models (***Yu et al., 2009***; ***Zhao and Park, 2017***; ***Keeley et al., 2019***), and nonlinear tuning curve models (***Zhang et al., 1998***; ***Zemel et al., 1998***; ***Cronin et al., 2010***; ***Rad and Paninski, 2011***; ***Calabrese et al., 2011***; ***Park et al., 2014***; ***Savin and Tkacik, 2016***; ***Rad et al., 2017***; ***Wu et al., 2017***).

Calcium imaging (CI) is a popular approach for recording large neural populations due to its ability to image large areas (>0.5mm^2^) at micron level resolution, enabling the simultaneous recording of many neurons (100-1000 typical, 10^6^ in advanced systems (***Demas et al., 2021***)) and tracking the same population of cells for days at a time. CI, however, measures neural spiking activity indirectly. Specifically, fluorescence changes in recorded CI time series represent fluctuations in intra-cellular calcium concentrations that result from the biophysics of action potentials; each spike results in a rapid rise in the calcium concentration. The fluorescence then jumps as the calcium is bound to the fluorescent proteins followed by a slower decay as the calcium unbinds (***Song et al., 2021***; ***Helmchen and Tank, 2015***). While theoretically the relationship between neural spiking and calcium-based fluorescence is well characterized, practically, variability in concentrations and noise considerations complicate the ability to discern single spikes (or even small bursts of 2-3 spikes) from calcium traces (***Song et al., 2021***; ***Ledochowitsch et al., 2019***).

Due to the indirect relationship between neural firing and CI traces, point-process models are no longer directly applicable to CI data. Instead methods to statistically analyze CI datasets commonly take two approaches: (1) ignore spiking and resort to Gaussian noise models operating directly on the calcium traces; or (2) apply Poisson models to *estimates* of the spike train obtained from calcium inference methods (***Smith and Häusser, 2010***; ***Ko et al., 2013***; ***Pnevmatikakis et al., 2016***). The former approach is suboptimal because the statistics of CI data are asymmetric with high-skew and long tails, and thus widely deviate from the statistics of Gaussian distributions (***Wei et al., 2020***). Additionally, *i*.*i*.*d*. noise observations are not appropriate for the long time-scale autocorrelations observed in CI data. The latter approach, spike-time estimation, is suboptimal because it does not take into account the uncertainty of the unobserved spikes. Specifically, single spikes have been shown to only be visible in the CI traces a fraction of the time (***Huang et al., 2021***), and the nonlinearities in calcium buffering in bursting can make spike estimation highly unreliable in many settings. Our approach instead uses an observation model that is more faithful to the data-generating process, which we demonstrate provides more accurate inference and scientific insight.

Specifically, here we extend a calcium observation likelihood first presented in ***Ganmor et al. (2016***) to latent variable models (LVMs) where the firing rate is a function of the latent variables. The three primary LVMs we consider are hidden Markov models (HMMs) (***Smith and Brown, 2003***; ***Escola et al., 2011***; ***Krause and Drugowitsch, 2022***), Gaussian Process Factor Analysis (GPFA) (***Yu et al., 2009***), and Latent Factor Analysis via Dynamical Systems (LFADS) (***Pandarinath et al., 2018***), though our approach also applies to a wide variety of models, including switching dynamical systems (***Linderman et al., 2017***) and more general nonlinear dynamical models (***Mudrik et al., 2024, 2025***; ***Yezerets et al., 2025***; ***Chen et al., 2024***). The LVMs we consider here are fit using diverse inference procedures—including maximum likelihood for HMMs and variational inference for GPFA and LFADS—highlighting the broad applicability of our framework. We evaluate the models using simulated datasets, a state-of-the-art biophysical simulator NAOMi (***Song et al., 2021***), and in-vivo 2-photon calcium imaging recordings. In each case, we demonstrate that the calcium likelihood, used in conjunction with various LVMs, recovers underlying latent structure more accurately than traditional observation models.

## Results

### Calcium LVM Framework

We propose the following framework (Fig. 1) for adapting LVMs developed for spiking data to the setting of calcium imaging recordings **y**_**t**_ ∈ ℝ^**N**^ of *N* neurons for *t* ∈ {1, …, *T*}. We consider models that prescribe a generative distribution over Poisson spike counts **s**_**t**_ ∈ ℕ_**0**_^**N**^ via latent variables **x**

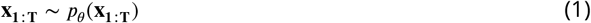

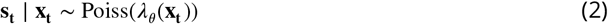

where the latent variables **x** may be continuous or discrete and the firing rate λ_θ_ depends on the value of the latent variables, potentially through a mapping parameterized by θ. This formulation includes many common LVMs used in neuroscience that we will describe in detail in the following sections. Previously, ***Ganmor et al. (2016***) proposed a model for fitting firing rates from calcium traces via approximate marginalization over unobserved Poisson spike counts. Here we propose to augment the generative models over spike counts with this model such that for a given neuron *n*

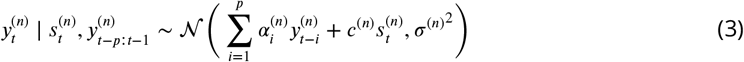

where we have generalized the approach of ***Ganmor et al. (2016***) to the setting of higher-order AR processes. This model has three sets of parameters per neuron. The AR coefficients α_*i*_ account for autoregressive calcium dynamics. If the model is *AR* (1), then α_1_ determines the exponential decay of fluorescence. Higher-order AR models (e.g., *AR*(2)) can additionally capture a finite rise time for the fluorescence response following a spike, rather than the instantaneous jump assumed under *AR*(1) (***Pnevmatikakis et al., 2016***). In practice, the order *p* can be chosen via model comparison (e.g., comparing held-out predictive likelihood across AR orders, as we do for GPFA below) or set according to the known kinetics of the calcium indicator used. Next, *c* describes the fluorescence increase due to a single spike and σ^2^ is the variance of additive Gaussian noise.

**Figure 1.**
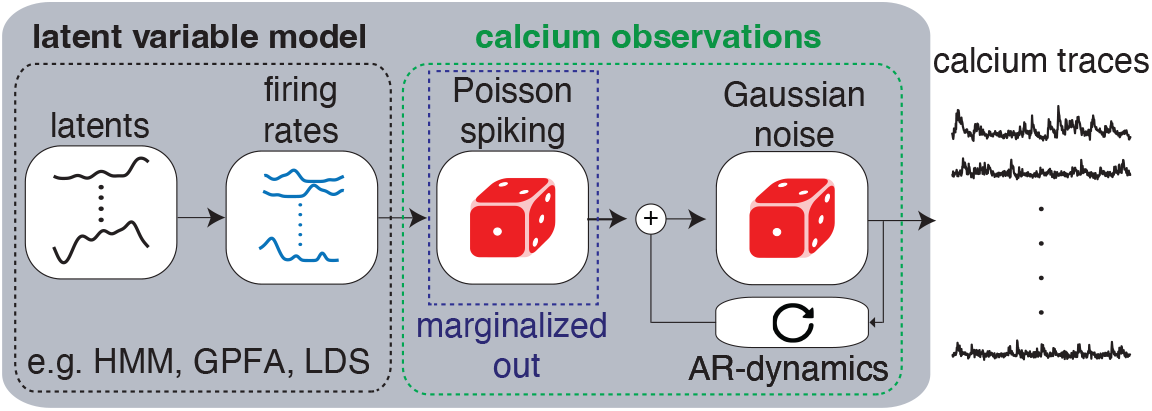
Schematic for the calcium LVMs. A low-dimensional latent variable maps to Poisson firing rates for time-series neural population activity. These Poisson spike probabilities are marginalized over and fed through an auto-regressive process to describe the evolution of the calcium traces.

Importantly, in this model the unobserved spike count variables can be approximately marginalized out via numerical integration such that it is tractable to evaluate 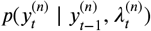. Accordingly, with automatic differentiation approaches and current approximate inference methods, this observation model can be generally applied across a variety of neural population models. In the following sections we will detail three specific applications of this framework.

### Calcium Hidden Markov Models

Hidden Markov Models (HMMs) are latent sequence models for identifying discrete structure over time. Here we describe the calcium HMM, an HMM with a more appropriate observation distribution for identifying discrete sequential structure in calcium imaging recordings. The generative model for a population of neurons is

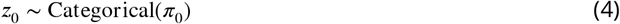

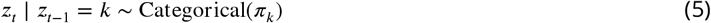

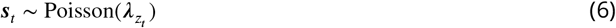

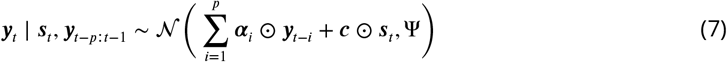

where *z* ∈ {1, …, *K*} is a discrete variable taking on one of *K* values, **y**_*t*_ ∈ ℝ^*N*^ and ***s***_*t*_ ∈ ℕ_0_^*N*^ are the vectors of fluorescence measurements and spike counts, respectively, across all *N* recorded neurons at time *t*, ⊙ indicates element wise multiplication, and Ψ is a diagonal matrix of per-neuron noise variances. The parameters π_0_ and π_*k*_ correspond to the initial discrete state distribution and transition distribution conditioned on state *k*, respectively. Here, we have rewritten the per-neuron observation model in Equation (3) in vectorized form; in the simulated and real-data experiments below we use an AR(1) process (i.e., *p* = 1), though the general AR(*p*) form above applies equally to higher-order dynamics. In the model, each discrete state prescribes a separate Poisson spike rate for every neuron. The spike counts generated from the model are then input to the AR observation process, whose parameters are shared across states.

The model is fit via the Expectation-Maximization (EM) algorithm (***Dempster et al., 1977***). The spike counts are numerically marginalized to get state-dependent likelihoods *p*(**y**_*t*_ ∣ **y**_*t*−1_, *z*_*t*_ = *k*) used in the E-step. In the M-step, the expected joint log likelihood is optimized with respect to the model parameters using automatic differentiation.

### Simulated Data

We demonstrate the potential benefits of the Calcium HMM in a simulated experiment where a population of neurons exhibits sequential firing activity (Fig. 2a-b). The data is simulated from an HMM with 5 states and a separate cluster of neurons has high firing rates in each state. The transition matrix enforces sequential transitions between clusters. We first simulated the spike counts and then simulated the calcium observations given the spike counts. The calcium observations were generated from Equation (7) with additional independent measurement noise per time point. Importantly, this additional noise is biophysically motivated, meaning that the data do not fully arise from any of the models we will consider and are more faithful to real data collected in neuroscience experiments.

**Figure 2.**
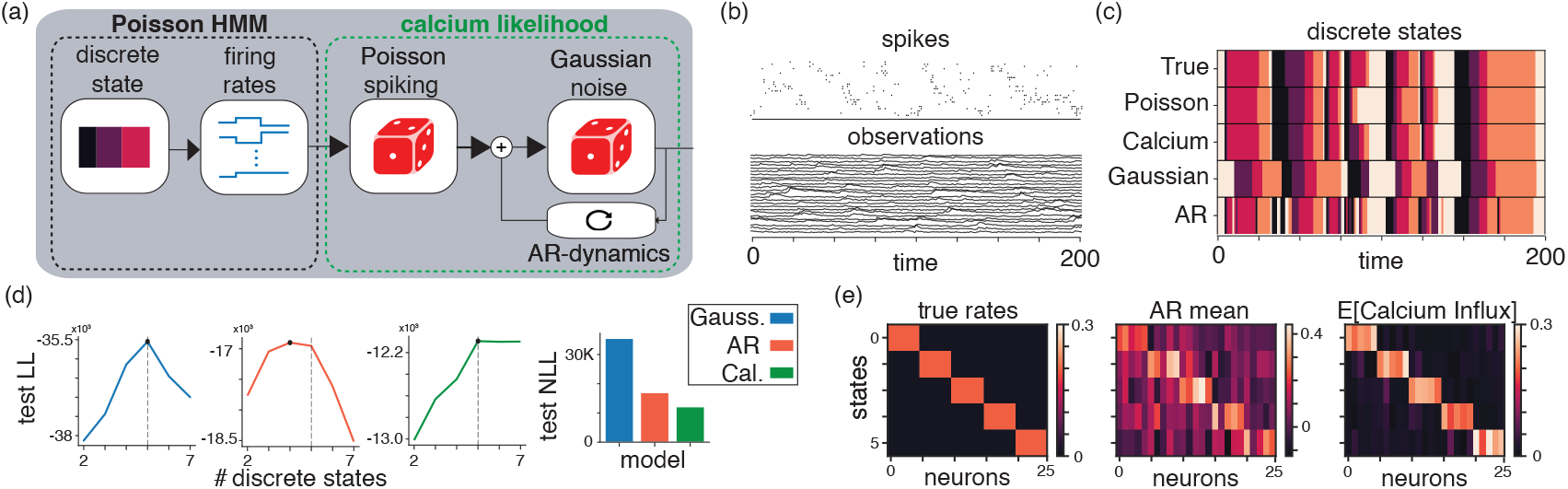
Simulated Calcium HMM. (a) Schematic for the overall Calcium HMM model. (b) The simulated data and (c) the inferred and true discrete states of the underlying latent states using different calcium models. (d) The inferred test log-likelihoods as a function of the number of latent discrete states for different observation models (left) and the resulting test negative log-likelihood across models given the ground truth number of discrete states (right). (e) True neural rates for each state of the underlying data (left) alongside the inferred AR parameter mean (middle) and expected calcium fluorescence value (right).

We fit four different models to the simulated data. First, we fit a Poisson HMM with five states to the underlying spike counts. This comparison point quantifies how informative the underlying spikes are for recovering the model parameters and latent states, providing a reasonable upper bound on performance. We next fit three HMMs with different observation models to the generated calcium imaging data. The three observation models are independent Gaussian, autoregressive (AR) Gaussian, and the calcium observation model from Equation (7). The AR Gaussian model can be thought of as a special case of the calcium model that ablates the spiking component to test the relative utility of including the discrete spiking.

We found that modeling the calcium imaging data as an autoregressive model with a latent discrete spiking component best explained the simulated data. On test data, the latent states recovered by the calcium HMM (*ρ* = 0.89) matched the ground truth far more accurately than those from the Gaussian HMM (*ρ* = 0.53) or autoregressive HMM (*ρ* = 0.73). Notably, the calcium HMM nearly matched the performance of the Poisson HMM (*ρ* = 0.91) fit directly to the true spiking data (Figure 2c). The calcium HMM achieved the highest test log-likelihood out of the models fit to the simulated calcium data and correctly identified the number of latent discrete states (Fig. 2d). Finally, for each discrete state we compared the true neuron firing rates with the “calcium influx” **c** ⊙ λ_**z**_, which are the firing rates inferred from the Calcium HMM model scaled by the influx parameters. We find that the estimated calcium influx approximately matches the true simulated firing rates and correctly identifies the different clusters of neurons, which is in contrast to the AR model that learns a less clear clustering (Fig. 2e).

### Modeling Piriform Cortex Recordings During Odor Presentation

After validating calcium HMM in simulation, we next demonstrate fitting the model to calcium imaging responses recorded in mouse piriform cortex during passive odor presentation (***Daste and Pierré, 2022***; ***Srinivasan et al., 2023***). In this dataset, ten different odors were presented eight times over the course of a session containing 80 trials. The duration of each trial was 30 seconds and the odor was passively presented for 1 second starting 10 s after the trial onset.

We fit HMMs with calcium or Gaussian observation models to the responses of 284 neurons in piriform cortex recorded via calcium imaging (Fig. 3a). We first selected the number of discrete states *K* for each model via cross-validation (60 training trials, 19 test trials; *K* = 13 for calcium and *K* = 12 for Gaussian). The test log likelihood for the calcium HMM is substantially higher than that of the Gaussian observation model due to the more expressive observation model (Fig. 3b).

**Figure 3.**
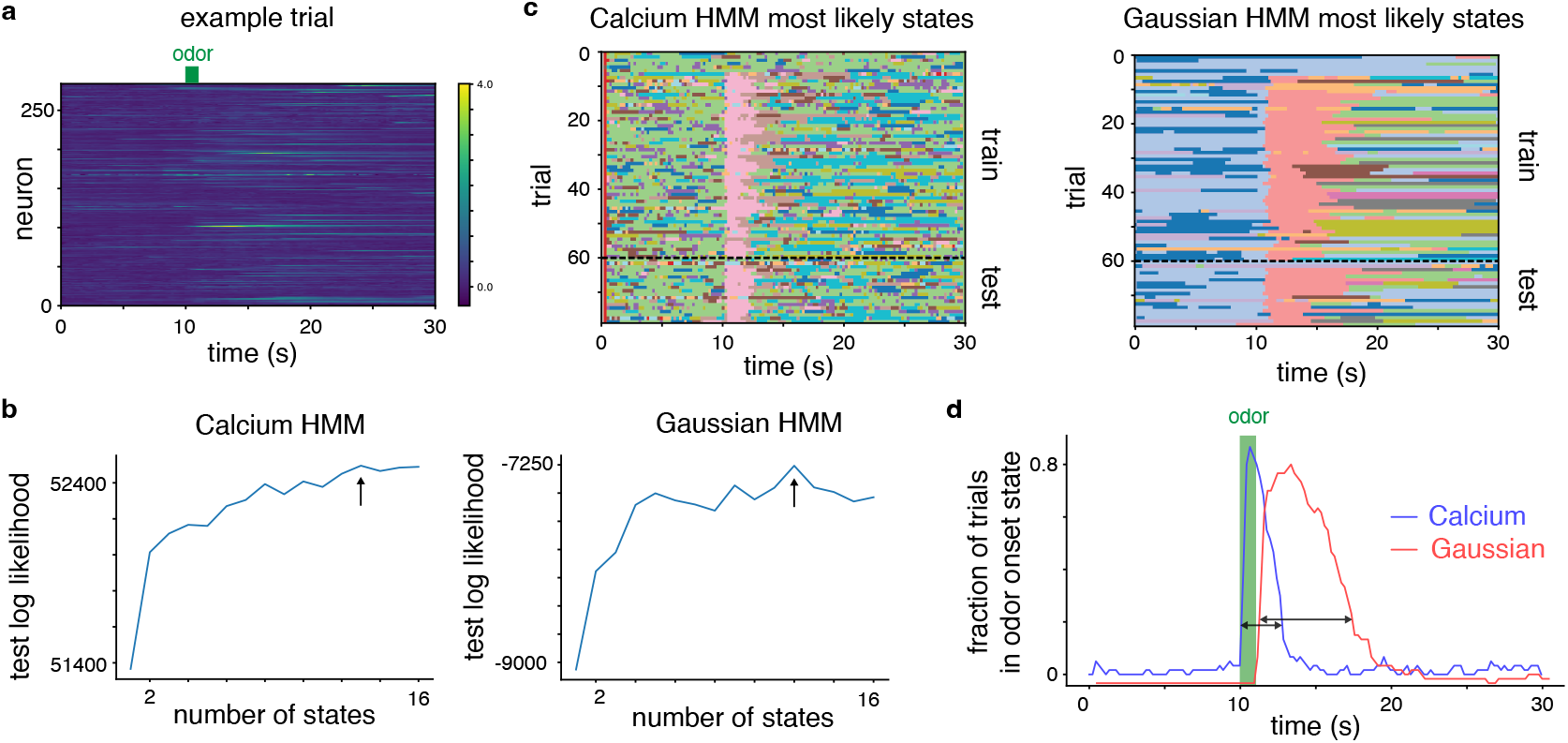
HMM comparison on odor response data. The calcium HMM identifies an odor onset state that is more tightly coupled with the actual odor onset.**(a)** Example Δ*F* /*F* traces for the population of recorded neurons on one trial. **(b)** Test log likelihoods for calcium and Gaussian HMMs as a function of the number of discrete states. Arrows indicate the number of discrete states with the highest test log likelihood. **(c)** Inferred most likely states on both training and test trials for each model. Each model identifies a consistent “odor onset” state linked to the time of odor presentation at 10s. **(d)** The fraction of trials in the odor onset state at each time point for each model. The calcium HMM odor onset state peaks more closely to the odor presentation window and has a shorter width (black arrows denote calculation of width).

We next analyzed the resulting calcium and Gaussian HMMs given the optimal number of states for each model. In each model, we found a discrete state that appears locked to odor onset across most trials (Fig. 3c). This “odor onset” state appears to mark a transition in the population response that occurs across most odor presentations. Notably, we found that the temporal extent of the odor onset state for the calcium HMM was more closely locked to the odor delivery window than the odor onset state for the Gaussian HMM (Fig. 3d). For the calcium HMM, trials generally transitioned into the odor onset state shortly after the odor presentation started and transitioned out of the odor onset state after a couple of seconds (peak onset state occupancy at 10.63s and onset state occupancy width of 2.65s). However, for the Gaussian HMM the transition into the odor onset state was generally delayed relative to the odor presentation and had a longer duration (peak onset state occupancy at 12.85s and onset state occupancy width of 6.64s).

The differences in the inferred odor onset state highlight the potential utility of the calcium LVMs described in this paper. In this dataset, the calcium HMM identified a discrete state that was time-locked to a behavioral variable of interest (odor presentation) in terms of both onset and duration of the discrete state. The calcium HMM odor onset state is more consistent with a population state that is time locked to the odor delivery window than the Gaussian HMM odor onset state. These results match our intuitions from the simulated calcium HMM example, where the inferred discrete states using the calcium HMM closely match the true discrete states but the inferred discrete states using the Gaussian HMM are generally delayed relative to the true discrete states.

### Calcium Gaussian Process Factor Analysis

Gaussian Process Factor Analysis (GPFA) is a standard tool for identifying smooth continuous latent structure underlying neural population data (***Yu et al., 2009***). Many modern variants of GPFA include count-observation likelihoods and are typically evaluated on spiking data (***Zhao and Park (2016***); ***Duncker and Sahani (2018***); ***Keeley et al. (2020b***); ***Wu et al. (2017***)). Here, we apply the calcium observation model to GPFA. In GPFA the prior on each latent dimension over time is a Gaussian Process. For a discrete set of time points, the prior is a multivariate Gaussian

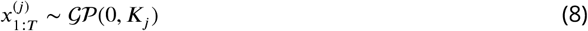

where the covariance *K*_*j*_ is a *T* × *T* matrix whose entries are defined by a covariance function *k*(*t, t*^′^). Here, we use the function *k*(*t, t*^′^) = exp(−(*t* − *t*^′^)^2^/(2*ℓ*^2^)) with a length-scale parameter *ℓ*. However, the proposed approach applies to other covariance functions as well. The dimensions are concatenated at each time point to produce a vector ***x***_*t*_ = [*x*^(1)^, … , *x*^(*p*)^]. The neural rates are generated via a linear map from the latent space followed by a non-linearity *f* which provides the rate parameter of the Poisson distribution. The observations are then modeled using the calcium AR process as before, however, for this example we include an additional AR-2 comparison, where the CI dynamics evolve dependent on the previous two time-steps.

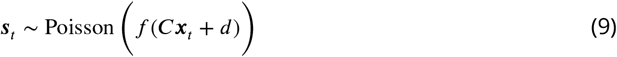

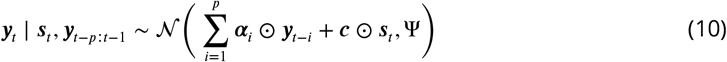

where *p* is set to either 1 or 2 for our GPFA comparisons. The linear transformation contains weights or “loadings” *C* and offsets *d*. Here, Ψ is a diagonal covariance matrix (Fig. 4b). The model is fit using a variational inference scheme which samples over the expectation term in the objective function (a so-called ‘black-box’ approach, see ***Keeley et al. (2020a***)).

**Figure 4.**
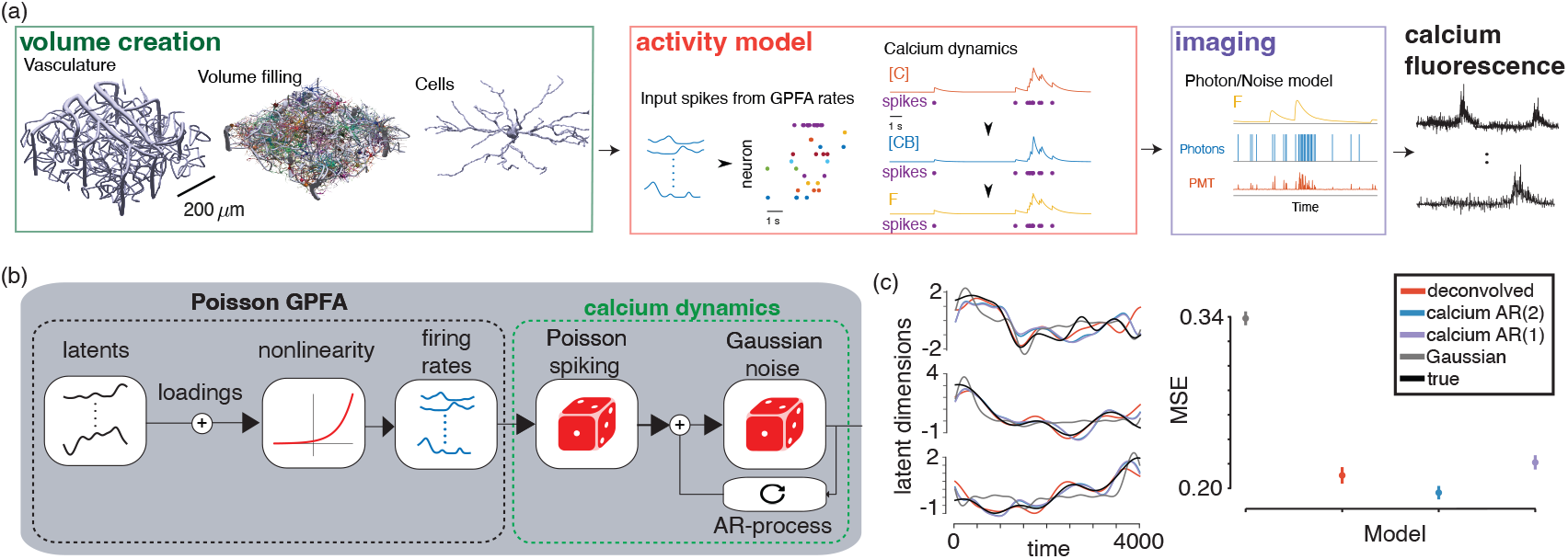
Calcium GPFA simulated experiment using biophysical calcium imaging simulator. (a) Graphical depiction of the biophysical calcium imaging simulator. (b) The Calcium GPFA model. (c) The temporal evolution of the three true underlying latent variables and the inferred latents from the population data using different observation likelihoods (left) and the overall estimation error of the latent variables under each model (right).

To evaluate the model, we used a biophysical calcium imaging simulator NAOMi (***Song et al., 2021***) to simulate a population of 30 neurons whose spiking activity was generated from a Poisson GPFA model with three continuous latent dimensions (Fig. 4a). The simulator was critical for obtaining a realistic recording of CI data with known *population* ground truth spiking. Specifically, NAOMi simulates data via the biological and optical processes underlying CI, including variability in expression, optical aberrations, and calcium dynamics, providing a fair comparison that is not simply data sampled from a similar statistical model used to fit the data. We fit GPFA models with CI and Gaussian observation models to the data.

The model with CI observations more accurately recovers both the continuous latent states (Fig. 4c,d), though in this example with the NAOMi-simulated traces, we find that the order of the AR process in the likelihood plays an important role in identifying the true underlying latent structure, with the CI AR2 likelihood outperforming AR1 as well as GPFA run on the deconvolved traces.

### Calcium LFADS

Latent Factor Analysis via Dynamical Systems (LFADS ***Sussillo et al. (2016***); ***Pandarinath et al. (2018***)) is a model of neural population spiking data with nonlinear, recurrent neural network (RNN) dynamics. The model is general and provides accurate fits to neural population activity in multiple real-world settings (***Pandarinath et al. (2018***); ***Pei et al. (2021***); ***Keshtkaran et al. (2022***)). As in the previous models, the standard LFADS formulation prescribes a distribution over Poisson firing rates via a Poisson GLM readout from the RNN. For calcium imaging data, previous work has proposed to incorporate an additional set of continuous latent variables corresponding to approximate spiking (***Prince et al., 2021***) or to fit the model to deconvolved spiking events (***Zhu et al., 2022***). Here we propose an alternative adaptation of LFADS to calcium imaging data by adding the calcium observation model on top of the firing rates. The schematic of the model extension can be seen in Figure 5a. Importantly, the standard amortized variational inference fitting procedure for LFADS (***Kingma and Welling (2014***); ***Kingma et al. (2015***); ***Sussillo et al. (2016***)) does not need to be modified with this change since the latent Poisson spike counts are marginalized out, in contrast to the model in ***Prince et al. (2021***). Additionally, the model is fit directly to fluorescence measurements without deconvolution, in contrast to ***Zhu et al. (2022***).

**Figure 5.**
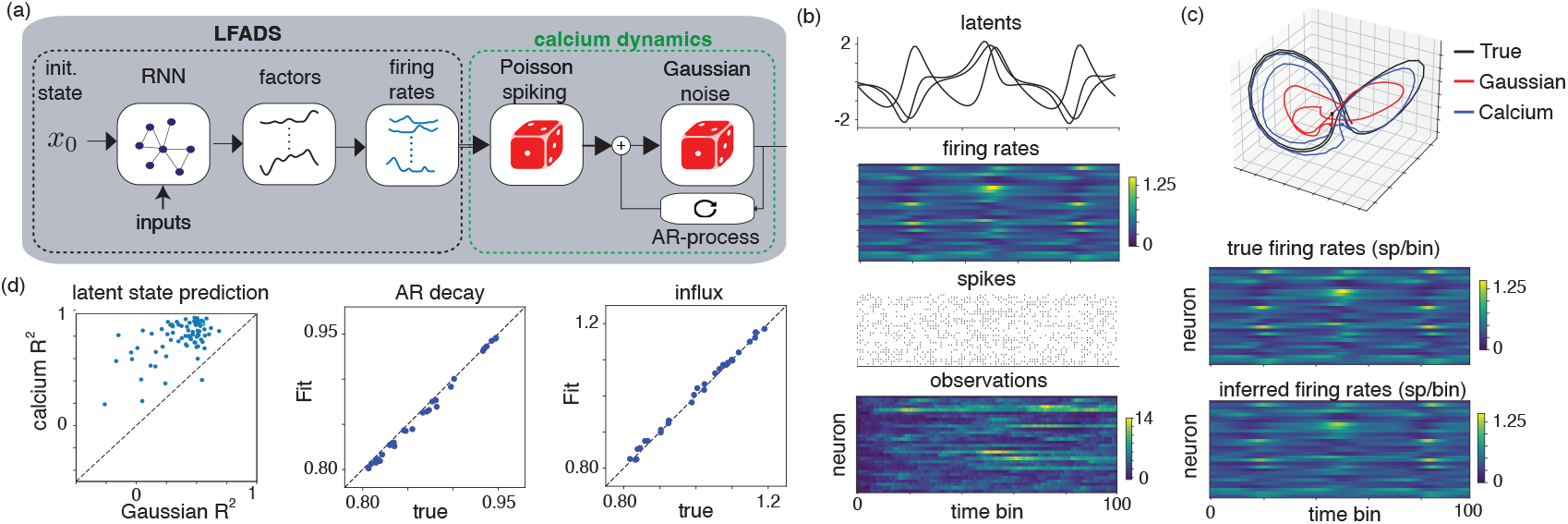
(a) Calcium LFADS model. (b) Generated latents variables as well as Poisson firing rates, spiking activity, and observed calcium traces. (c) inferred latent dynamics under an (AR1) calcium model and Gaussian likelihood. (d) latent state prediction performance as well as inferred and true calcium trace hyperparameters.

To demonstrate the approach, we synthetically generated latent time series from a 3D Lorenz attractor. We mapped the latent time series to Poisson spiking rates via a Poisson GLM. Then, we simulated Poisson spiking followed by autoregressive calcium dynamics (Fig. 5b). We compare the model with LFADS fit using a Gaussian observation model. In Figure 5c,d we show that the model with calcium observations infers more accurate latent variables than the model with Gaussian observations (mean latent state reconstruction *R*^2^ = 0.77 for calcium model compared to *R*^2^ = 0.37 for the Gaussian model, with 78/80 trials better reconstructed by the calcium model). Additionally, the inferred parameters of the calcium autoregressive dynamics are qualitatively similar to the ground truth parameters across the simulated neurons (Fig. 5d).

## Discussion

Here we have demonstrated that a tractable likelihood for calcium imaging data can be used to adapt a variety of latent variable models in neuroscience to the setting of calcium imaging recordings. The proposed models can be fit with similar computational and inference requirements to the equivalent spiking versions of the models and do not require deconvolution methods.

Our proposed method carries a few distinct practical dependencies. Our work depends on extracting the calcium traces from fluorescence videos, which is itself an active area of research with a number of available methods (***Pnevmatikakis et al., 2016***; ***Charles et al., 2022***; ***Pachitariu et al., 2016***; ***Dinç et al., 2021***). As such, errors in calcium imaging source extraction, e.g., due to false transients (***Gauthier et al., 2022***), can impact the outputs of our model in much the same way that multi-unit recordings affect electrophysiology analysis. Care, therefore, should be taken to curate or validate the calcium traces, or use a robust estimation method (***Gauthier et al., 2022***; ***Inan et al., 2017***).

Methodologically, our work complements a variety of recent efforts for improved estimation of neural activity from calcium imaging data. Many of these approaches work with the implicit goal of best inferring spikes from calcium traces (***Pachitariu et al., 2018***; ***Wei et al., 2020***). Although such efforts are useful, they do not take into account the uncertainty underlying spiking in estimating neural rates.

In contrast to these two-step spike-estimation approaches, there has been recent work that also uses generative models to describe calcium dynamics themselves, avoiding the need to first deconvolve and then infer latent structure from spike trains. In particular, ***Prince et al. (2021***) uses a variational auto-encoder-style model to directly infer latent dynamics from raw calcium traces using latent Poisson rates and observed Gaussian likelihoods on the resultant calcium traces. Additionally, ***Koh et al. (2022***) derives a generative autoregressive model of calcium dynamics from underlying latent variables that evolve via linear dynamics. Here, there is no spiking explicitly modeled, and the underlying latent dynamics directly prescribe correlations in the observed calcium traces. Similarly, ***Rupasinghe et al. (2021***) uses signal and noise correlations in the calcium population activity to extract latent time-series which generate latent (Bernoulli) spike rates, which, similar to our approach, leverage uncertainty in latent spike counts. This again avoids the two-step procedure and directly infers latent structure from population covariance.

Our work here as well as in ***Ganmor et al. (2016***) complements these approaches, but in contrast to them, does not enforce a specific model of latent structure. Instead, we have shown how a versatile calcium likelihood can be used in conjunction with a wide variety of latent variable models used in neuroscience. While we demonstrate the effectiveness of this approach for GPFA, LFADS, and HMMs, there are many other existing latent variable models that could be adapted for our approach including switching dynamical systems (***Linderman et al., 2016b***; ***Zoltowski et al., 2020***; ***Karniol-Tambour et al., 2022***), other nonstationary dynamical systems (***Mudrik et al., 2024, 2025***), extensions to GPFA (***Keeley et al., 2020a***; ***Gokcen, 2023***), or other auto-encoder style latent variable models (***Zhou and Wei, 2020***; ***Schneider et al., 2023***). Furthermore, because our framework maintains the raw fluorescence as a fixed observation, it enables principled model comparison—such as computing cross-validated predictive likelihoods on held-out neurons—across any of these varied latent architectures. Although benchmarking of specific models is beyond the scope of this study, our approach provides the statistical infrastructure necessary for practitioners to evaluate and select the most appropriate latent model for their data.

Beyond traditional latent variable frameworks, our proposed approach may also be relevant for large-scale foundation models of neural recordings (***Azabou et al., 2023***; ***Ye et al., 2023***; ***Zhang et al., 2024***). Such models often incorporate Poisson spiking rates, and their reconstruction losses typically use Poisson or cross-entropy observation models. The CI observation model could be applied to directly adapt such models for CI data and even presents the opportunity to jointly train large-scale models on electrophysiological and CI recordings.

While we focus in this work on calcium imaging, this plug-and-play likelihood model can be more generally applicable to any indicator that has an underlying point process observed through an exponential rise/decay dynamic. Specifically, all sensors (aside from some voltage sensors) have such a rise and decay time and thus the limiting factor in using the model more broadly is the biological target of the sensor. For example, glutamate (***Marvin et al. (2013***)), dopamine (***Patriarchi et al. (2018***)), and similar indicators all can be considered to be driven by the action potentials, i.e., the Poisson-like firing model. Other sensors, however, are reflective of other biological processes, such as hemodynamics (via imaging, see ***Glover (2011***)) or ultrasound (***Brunner et al. (2021***)) or neuromodulation (Norepinephrine imaging with nLight as in ***Rohner et al. (2026***)). These processes are smoother and more continuously varying and therefore would require replacing the Poisson dynamics with an appropriate alternative. Finally, voltage imaging (***Knöpfel and Song (2019***)) is often fast enough to resolve single spikes, removing the necessity of the hierarchical model and enabling point-process models to be used directly.

Overall, we demonstrate that models developed for direct use on neural spike-trains can be adapted to calcium imaging data using a simple plug-and-play approach. Our public repository^1^ is available in both JAX and PyTorch implementations, with tutorials to demonstrate how they can be integrated into existing models, and it is our hope that this method will accelerate the application of powerful latent variable models to calcium imaging data.

## Methods

### Calcium observation model

A likelihood originally proposed in 2016 (***Ganmor et al., 2016***) defines a conditional probability of measured calcium fluorescence *y* (the so-called “dF over F”) given a Poisson firing rate λ. This model is an autoregressive (AR) model whose output depends linearly on its previous value one timestep in the past (so-called AR(1) model). This basic AR(1) version additionally has the calcium level depend on Poisson spiking with additive independent Gaussian noise on each time step:

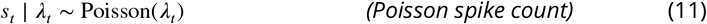

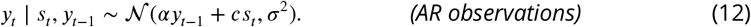

This model has three parameters:

- α, the AR coefficient, which determines exponential decay of fluorescence;
- *c*, the fluorescence increase due to a single spike;
- σ^2^, the variance of the additive Gaussian noise.

Practically, this model can be interpreted as a process where an individual spike causes an instantaneous rise in calcium florescence, followed by an exponential decay due to the AR coefficient. Here, additive Gaussian noise is fed through the AR process. Importantly, the model allows us to marginalize over spike counts *s*, so we can consider the probability of the fluorescence given the *rate*, λ, and we need not consider individual spikes. We elaborate on the details of the model below.

### Conditional distribution given spikes

To emphasize AR structure (shown here of order 1), we can rewrite the model with *y*_*t*_ terms on the left as follows:

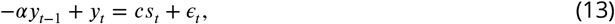

for time bins *t* = {1, … , *T*}, and we will assume that *y*_0_ = 0. This allows us to rewrite the model in vector form as Gaussian conditioned on the spike counts:

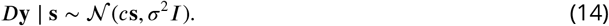

where **y** = (*y*_1_, … , *y*_*T*_)^⊤^ is the vector of fluorescence measurements, **s** = (*s*_1_, … , *s*_*T*_)^⊤^ is the vector of spike counts, and *D* is a matrix that computes weighted first-order differences:

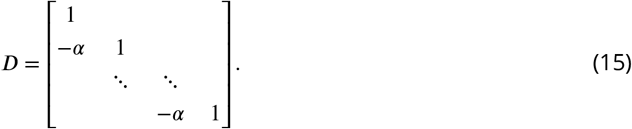

If we define a new variable δ = *D***y**, equal to the AR differences of the raw measurement data, we can write the conditional distribution as a product of independent terms

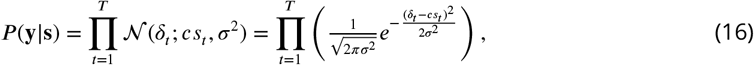

where δ_*t*_ = *y*_*t*_ − α*y*_*t*−1_ for the AR(1) case described above. This expression generalizes directly to higher-order AR processes; for a general AR(*p*) process, δ_*t*_ is given by

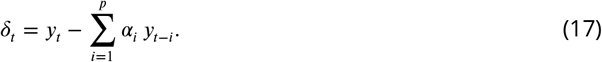

### Likelihood Evaluation

For model fitting and inference, we must be able to efficiently evaluate the likelihood of observed calcium responses marginalized over the unobserved spike count vector **s**. The independence across time bins in (16) means that these marginals can be computed independently for each time bin. That is, we can compute likelihood by summing over spike counts in each bin from 0 to some maximum possible spike count *R*.

Thus, numerical evaluation of the likelihood can be achieved practically as:

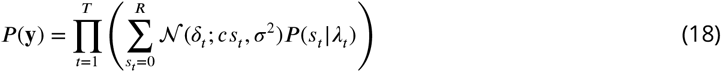

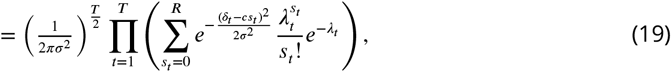

where 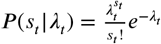 is the Poisson probability over spike counts in each time bin. In practice we of course use the log-likelihood, given by:

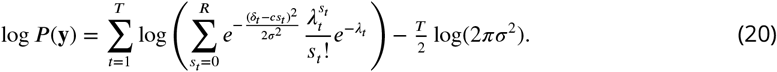

Importantly, this marginalization is amenable to automatic differentiation packages, making this a ‘plug-and-play’ observation distribution for various models.

### Synthetic HMM Dataset and Experimental Details

The synthetic HMM dataset had *K* = 5 states and *D* = 25 observed neurons. We generated two sequences of length *T* = 2000; the first was observed for training and the second was held-out for testing. The state transition matrix was designed to generate a repeating chain structure in the latent states such that the transition matrix *P* was

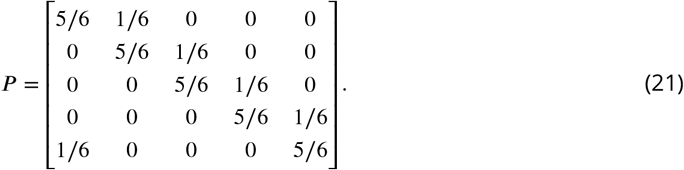

The synthetic neurons were grouped into 5 groups of 5 neurons corresponding to each of the latent states. Each neuron’s firing rate was 0.2 spikes per bin. The calcium observation model parameters were selected as α ∼ *N* (0.8, 0.1), *c* = 1.0, and σ^2^ = 10^−2^. The AR coefficients were clipped to be within the range α ∈ [0.6, 0.95]. Importantly, we added simulated measurement noise to the simulated calcium traces via a Gaussian noise model with zero mean and standard deviation of 0.2. Therefore, the simulated dataset is generated from a model that is mismatched to all models considered.

We initialized the calcium AR parameters via the following procedure. The AR coefficients were initialized via a linear regression predicting the next calcium observation from the previous observation for each neuron. The initial variance was set to the squared residual error of this linear regression. The initial fluorescence increases *c* were set randomly from the distribution *N* (1.0, 0.2).

### Piriform Cortex Recordings During Odor Presentation

We applied the Calcium HMM model to a publicly available dataset of piriform cortex calcium imaging recordings during passive odor presentation (***Daste and Pierré, 2022***). This dataset consists of 8 repeated presentations of 10 different odors, yielding 80 total trials. We used publicly available code to process the data into Δ*F* /*F* (https://gitlab.com/fleischmann-lab/datasets/daste-odor-set-2021-11). We additionally filtered out one anomalous trial and one anomalous neuron. After processing, the dataset consisted of recordings of 284 neurons across 79 trials. Each trial lasted 30 seconds and the imaging sampling frequency was 4.53 Hz (see (***Srinivasan et al., 2023***), ‘Two-photon microscopy’ methods for additional details).

We used the first 30 and last 30 trials as training trials and tested on the middle 19 trials. The observation model parameters were initialized with a two-step procedure. First, initial discrete state sequences were set using the known odor sequence or using K-means clustering. Then, we optimized the observation model parameters (both Poisson rates and calcium observation parameters) by maximizing the likelihood of the calcium observations with the discrete state sequence fixed. After this initialization, we then optimized all parameters with respective to the HMM marginal likelihood using SGD for 5000 iterations.

The calcium HMM was implemented in JAX using HMM tools from the Dynamax repository^2^. This was important for computational efficiency, as the optimization step was automatically batched across trials and compiled for speed.

### GPFA Synthetic Dataset with NAOMi Simulator

To use the NAOMi simulator, we generated a neural volume using the anatomy module and simulated the light propagation using the optics module. Rather than use the built-in Hawkes process to simulate arbitrary dynamics, we imposed the spike times from the GPFA spike generation using calcium dynamics module’s feature that enables user-defined spikes. The calcium module then generated the ground-truth fluorescence traces from the provided spikes and we generated the CI simulated videos using the scanning module.

To recover single fluorescence traces from the video we used profile-assisted least-squares (PALS) as in the original NAOMi paper (***Song et al., 2021***). In this process we scan the volume under no-noise conditions with only one neuron “on” at a time and no neuropil or other contamination. These scans provide the ground-truth spatial profiles that can be used to identify the temporal profiles by solving a per-frame least-squares optimization. To reduce noise, we regularize the time-traces with a sparsity-promoting *l*_1_ norm, also as in (***Song et al., 2021***). The PALS traces remove the confounding factor of cell detection, which can vary significantly between approaches and can result in bleed-through errors that can affect the inferred coding properties of the cells (***Gauthier et al., 2022***).

To generate the simulated data for Calcium GPFA inference, we generated Poisson spiking activity from 30 neurons for 4000 timepoints derived from a 3-dimensional latent space, where each latent is governed by a Gaussian Process with a different temporal length scale *ℓ* = {250, 450, 500}. Spike-times from these 30 neurons were then used as the ground-truth spikes in the NAOMi simulator. However, after simulation only 22 calcium traces had nonzero calcium dynamics. Therefore, inference for calcium GPFA was used on 22 calcium traces. The competing models either used Gaussian observations or Poisson observations from spikes that were determined by deconvolution via SpikeML (***Deneux et al., 2016***). GP length scales were all initialized to 350, calcium AR1 and AR2 parameters were initialized to 0.51 and {1.81, −0.81} for all neurons, respectively. The noise parameter per neuron was initialized by setting it to the variance of the calcium values determined across consecutive timepoints, and the amplitude of the calcium influx due to a spike was initialized to 1 for all neurons. The model was learned via black-box variational inference for 20000 iterations. All inferred latents were regressed to the true latent before calculating the mean squared error.

### Calcium LFADS Model and Experimental Details

To evaluate the Calcium LFADS model, we simulated synthetic calcium observations from the Lorenz dynamical system across 400 trials each of length 100 with Δ_*t*_ = 0.025 using the Runge-Kutta method (RK4). We simulated observations from 30 neurons. The average number of spikes per bin was 0.42. The calcium observation model parameters for each neuron were randomly sampled with α ∼ Unif(0.8, 0.95) and *c* ∼ Unif(0.8, 1.2). The autoregressive dynamics noise was set to 10^−3^. After simulating the calcium traces, we added zero-mean Gaussian measurement noise with a standard deviation of 0.2 independently to each time step. Importantly, this measurement noise is not present in the generative model, as in the HMM example.

The full generative model of the LFADS model with calcium observations is

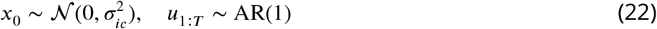

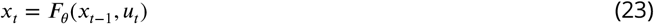

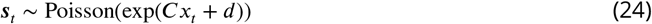

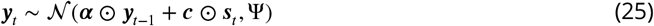

where *F* is a recurrent neural network (GRU) and the random inputs *u*_1:*T*_ are generated from an autoregressive process with *u*_*t*_ ∼ *N* (*au*_*t*−1_ + *b*, σ^2^). The model is fit as a sequential variational autoencoder as in (***Pandarinath et al., 2018***) with encoder networks inferring approximate posterior distributions over the initial state *x*_0_ and sequence of random inputs *u*_1:*T*_ . Before fitting the model, we initialized the calcium observation model hyperparameters α and *c*. For each neuron, the AR parameter α was estimated via a linear regression predicting *y*_*t*_ from *y*_*t*−1_ for timepoints where *y*_*t*−1_ > *y*_*t*_. The influx parameter *c* was estimated by sweeping over a range of possible values from 0.8 to 2.2 and identifying the value that best aligned the differences *y*_*t*−1_ − *y*_*t*_ with quantized values 0, *c*, 2*c*, 3*c*.

## Acknowledgments

SK was supported by the NIH BRAIN Initiative (F32MH115445-03). DMZ was funded by the Wu Tsai Interdisciplinary Postdoctoral Research Fellowship. ASC was supported by the NSF under Grant No. 2340338 (Faculty Early Career Development Program - CAREER). JWP was supported by the Simons Collaboration on the Global Brain (SCGB AWD543027), the NIH BRAIN Initiative (9R01DA056404), and a U19 NIH-NINDS BRAIN Initiative Award (U19NS104648).

## Footnotes

1 https://github.com/skeeley/Calcium_likelihoods

2 https://github.com/probml/dynamax

